# The aging mouse lipidome

**DOI:** 10.64898/2026.02.02.703178

**Authors:** Edgar Esparza, Steven E. Pilley, Xuanyi Shi, Preeti Iyengar, Shruti Sivaraman, George Wang, Racheal Mulondo, Sriraksha Bharadwaj Kashyap, Darius Moaddeli, Aeowynn J. Coakley, Ryo Higuchi-Sanabria, Whitaker Cohn, Peter J. Mullen

**Affiliations:** Department of Immunology and Immune Therapeutics, Keck School of Medicine, University of Southern California, Los Angeles, CA, USA; Alfred E. Mann School of Pharmacy and Pharmaceutical Sciences, University of Southern California, Los Angeles, CA, USA; Norris Comprehensive Cancer Center, Keck School of Medicine, University of Southern California, Los Angeles, CA, USA; Leonard Davis School of Gerontology, University of Southern California, Los Angeles, CA, USA

**Author notes:** Senior author (Peter J. Mullen).

## Abstract

Aging is associated with widespread metabolic changes that contribute to functional decline and disease. While prior studies have characterized age-associated changes in lipids, it still remains incompletely understood how the lipidome changes across tissues and between sexes during aging. Here, we performed targeted lipidomics across 10 organs collected from male and female mice at five ages spanning adolescence to old age. We analyzed 775 lipids across multiple lipid classes and found that aging affects the lipidome in an organ-specific manner. The thymus and quadriceps muscle had the most age-associated lipid changes, whereas lipid levels in organs such as the kidney and lung remained more stable. In quadriceps muscle, aging was associated with a decrease in specific phosphatidylcholine and phosphatidylethanolamine lipids, particularly those containing adrenic acid. We also identified sex-dependent differences in lipid composition, with the spleen showing differences throughout life. Spleens from female mice had lower levels of lysophosphatidylcholine and lysophosphatidylethanolamine compared to males. Together, these data provide a comprehensive atlas of age- and sex-associated lipid changes across mouse organs and complement existing metabolic and transcriptomic resources to support studies of mouse aging.

## Introduction

Humans are living longer, with the average human lifespan increasing from 66.8 years in 2000 to 73.4 years in 2019 (World Health Organization). But this has also been accompanied by a rise in the incidence of age-related diseases, including cancer, neurodegenerative diseases, and cardiovascular disease^1^. This trend underscores the critical need to improve our understanding of the aging process to limit the burden of these diseases and promote healthy longevity. Aging is accompanied by multiple metabolic changes^2^ and many successful anti-aging therapeutic strategies have targeted metabolism, highlighting the intersection between metabolic changes and longevity. Methionine restriction has been shown to increase longevity across multiple model organisms, including Drosophila^3^, rats^4^, and mice^5^ as well as improved neuromuscular function in 5xFAD mice, an aggressive model of Alzheimer’s disease^6^. Dietary supplementation with α-ketoglutarate (AKG) has also emerged as a promising anti-aging intervention, extending lifespan in *C. elegans*^7^ and in mice^8^, while also ameliorating the progression of osteoporosis in aged mice by promoting the osteogenic potential of bone marrow mesenchymal stem cells (MSCs)^9^.

In addition to polar metabolites, lipids play important roles in the aging process, and targeting lipid metabolism has been suggested as a strategy for extending lifespan and mitigating age-related decline^10^. In *C. elegans*, dietary supplementation with omega-6 polyunsaturated fatty acids arachidonic acid and di-homo-gamma-linoleic acid, as well as alpha-linoleic acid, promoted resistance to starvation and extended lifespan^11^. In mice, dietary supplementation with steroid 17-a-estradiol significantly increased lifespan in a sex dependent manner, increasing the male median lifespan by 12%^12^. Removal of visceral fat from rats significantly increases median and mean lifespan and can delay the onset of diabetes^13^. Genetically modifying lipid related genes has also been a strategy for increasing lifespan. Disrupting levels of the triglyceride synthesis enzyme acyl-CoA:diacylglycerol acyltransferase 1 (DGAT1) increased lifespan in female mice and inhibited the age induced increase of body fat and inflammation in white adipose tissue^14^. Together, these findings demonstrate that lipid based interventions are critical not only for extending lifespan, but also for reversing or inhibiting age-related phenotypes to promote healthy aging across multiple model organisms.

Our previous work characterized age- and sex-associated changes in polar metabolites across multiple mouse organs^15^. However, this analysis focused exclusively on polar metabolites and did not examine lipids, leaving a gap in our understanding of how metabolism as a whole changes with aging.

Previous studies have taken multiple different approaches to profile how the lipidome changes throughout life. One study performed longitudinal lipidomic profiling of human plasma from over 1,500 individuals, with some participants followed for up to nine years^16^. This identified lipid changes associated with insulin resistance and aging, suggesting important roles for lipids in immune and inflammatory regulation. Most lipid classes increased with age in the plasma, with prominent increases in ceramides, sphingomyelins, and cholesterol esters. However the study did not investigate how organ lipidomes change with age. Another study profiled 11 mammalian species (including humans) ranging from 3 to 120 years and found that the plasma lipidome could be used to predict the animal’s lifespan, with plasma long chain free fatty acids being inversely correlated with longevity^17^. In the longest living mammal (bowhead whale) and the longest living rodent (naked mole rat), changes in the lipidome are associated with longevity and their ability to avoid age-related deficiencies.^18,19^ emphasizing the need to further characterize the metabolic changes associated with aging.

It is also important to understand organ-specific changes in the lipidome, as plasma lipids may not fully represent changes at the organ level, as shown in our metabolic atlas of mouse aging. A previous study profiled changes in lipids across 13 organs in germ-free and specific pathogen-free mice at multiple ages (2, 12, 19, and 24 months) and found that aging was associated with increased bis(monoacylglycero)phosphate lipids containing polyunsaturated fatty acids, alongside decreased phospholipids enriched in saturated and monounsaturated fatty acids^20^. Increased BMP during aging was seen in a separate study comparing changes in lipids between 3-month-old and 24-month-old mice^21^. Another study characterized the mouse brain lipidome using CNS-derived primary cell cultures and tissue samples from distinct brain regions to identify cell type- and region-specific differences in lipid composition^22^. Here, mice of different ages and dietary conditions were analyzed to assess age- and diet-induced changes in brain lipid composition. Diets enriched in n-3 fatty acids, including eicosapentaenoic acid (EPA) and docosahexaenoic acid (DHA), led to significant alterations in phospholipid fatty acid composition. Additionally, age-associated differences were observed in HexCer and PE O-species are consistent with altered myelination and lipid metabolism in the aging brain^23^. Together, these studies show that age-associated lipidome changes occur during aging, but they also remain incompletely characterized across tissues. This highlights the need for a comprehensive analysis of age-related changes in lipids in in microbiome-intact mice at multiple ages in multiple organs, that can be integrated with existing metabolic and transcriptomic datasets.

In this study, we describe how the mouse lipidome changes across 10 organs at five ages in both male and female C57BL/6 mice. We found that aging affects each organ differently, with some organs undergoing extensive lipidomic remodeling, while others remain relatively stable. We also observed that sex shapes the lipidome of specific organs. This work was performed in the same context as our previous metabolic atlas of mouse aging and the Tabula Muris Senis project^24^, enabling integration of transcriptomic, metabolic, and lipidomic data to provide a more comprehensive understanding of the effects of aging and sex across mouse organs, and to serve as a resource for studies investigating age- and sex-specific processes and disease.

## Results

### Overview of the data

To characterize age-associated changes in lipid composition across the mouse lifespan, we performed targeted lipidomics on 10 organs collected from mice at five time points: 1 month (adolescent), 3 months (young adult), and 18, 21, and 24 months (old age). For each age group, we analyzed six male and six female mice, enabling the simultaneous assessment of age- and sex-dependent changes in the lipidome. Immediately following anesthesia, we collected plasma, heart, thymus, lung, liver, spleen, pancreas, kidney, quadriceps muscle and brain, as per our metabolic atlas of mouse aging^15^ (Fig.1a).

**Figure 1:**
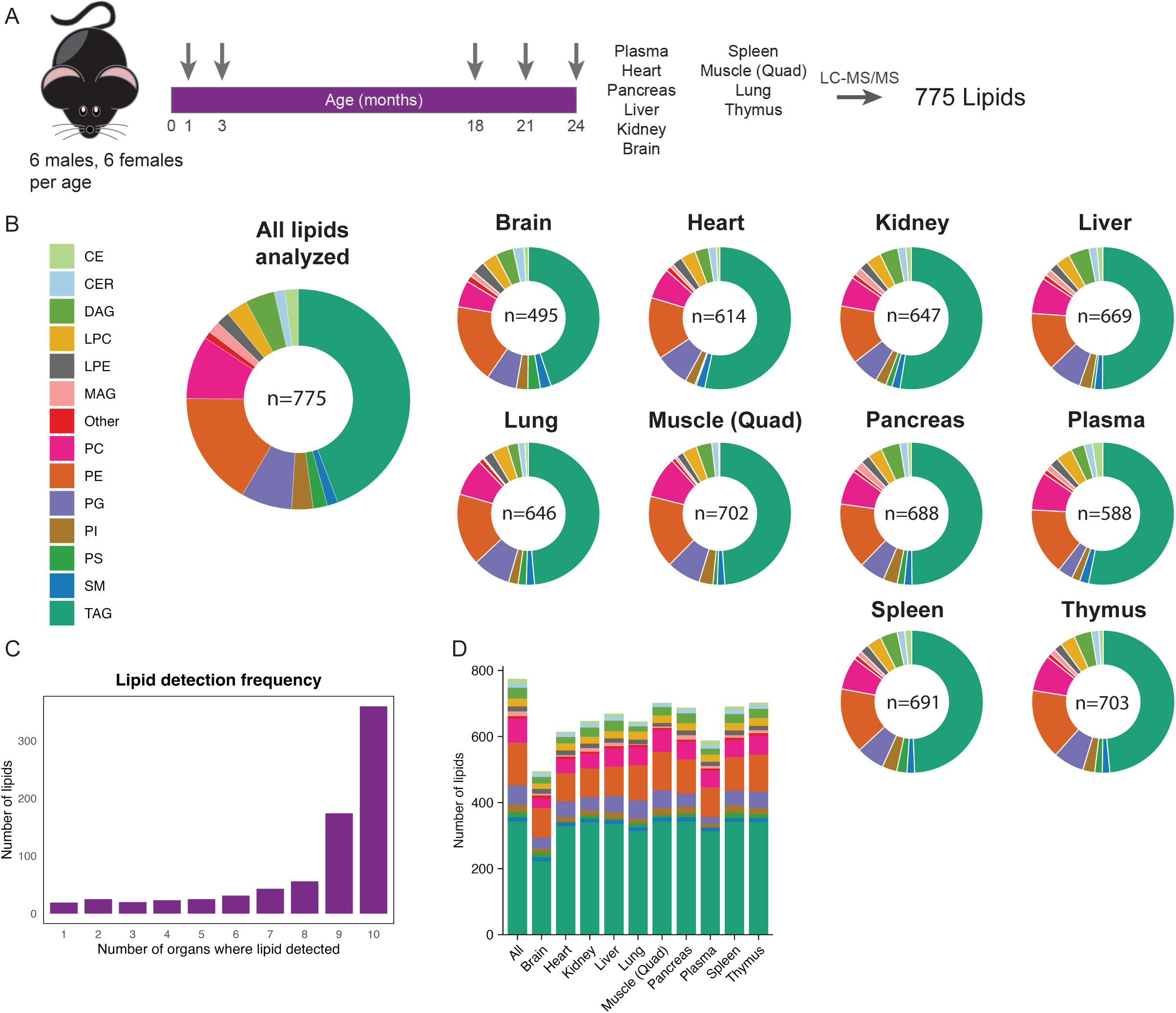
Overview of the lipid atlas of aging. **a**, Schematic of the experimental design. The arrows represent the ages at harvest. **b**, Distribution of lipid classes identified in (left) our targeted analysis and (right) each organ. **c**, The number of organs in which a lipid is found. **d**, Number of lipids of each class identified in each organ.

We extracted lipids using a methyl tert-butyl ether (MTBE)–based protocol and quantified lipid species by liquid chromatography coupled to tandem mass spectrometry (LC–MS/MS), using hydrophilic interaction liquid chromatography (HILIC) and multiple reaction monitoring (MRM) in positive and negative modes. We processed chromatograms in Skyline, identified lipid species based on curated fragment ions and retention times, and then normalized data to total ion counts and applied log transformation before downstream analyses.

Across all samples, we quantified 775 lipid species spanning multiple lipid classes, including triacylglycerides (TAGs), diacylglycerides (DAGs), monoacylglycerides (MAGs), phosphatidylethanolamine (PE), phosphatidylcholine (PC), lysophosphatidylethanolamine (LPE), lysophosphatidylcholine (LPC), ceramides (CER), phosphatidylinositol (PI), cholesterol esters (CE), phosphatidylglycerols (PG), phosphatidylserines (PS), and sphingomyelins (SM). TAGs were the most abundant lipid class detected, followed by PE and PC species. The thymus and quadriceps muscle had the greatest lipid diversity (703 and 702 detected lipids, respectively), whereas the brain contained the fewest (Fig. 1b).

Approximately half of all detected lipid species were present across all 10 organs, whereas a small subset of lipids were only identified in single tissues (Fig.1c). Lipid class composition was broadly conserved across organs, with the notable exception of the brain, which had fewer triglycerides compared to all other tissues (Fig.1d), consistent with previous studies showing that the brain lipidome is mostly comprised of phospholipids and sphingomyelins^22,25^. Together, these results demonstrate that while the proportion of each lipid class is similar between organs (Fig. 1b, 1d), there are organ-specific differences in the individual lipids found abundance (Fig. 1c).

### Organs have distinct lipid profiles

We next performed Spearman correlation analyses to assess similarities and differences in lipid composition across organs. Samples from the same organ correlated and clustered with each other (Fig. 2a), matching what we observed with polar metabolites^15^. The pancreas (0.69) and spleen (0.7) had the lowest intra-organ correlations suggesting variability in lipid composition across age and/or sex (Extended Data Fig. 1a). In contrast, the brain (0.97) and plasma (0.93) had the highest intra-organ correlations, indicating that lipid type was not as affected by age or sex.

**Figure 2:**
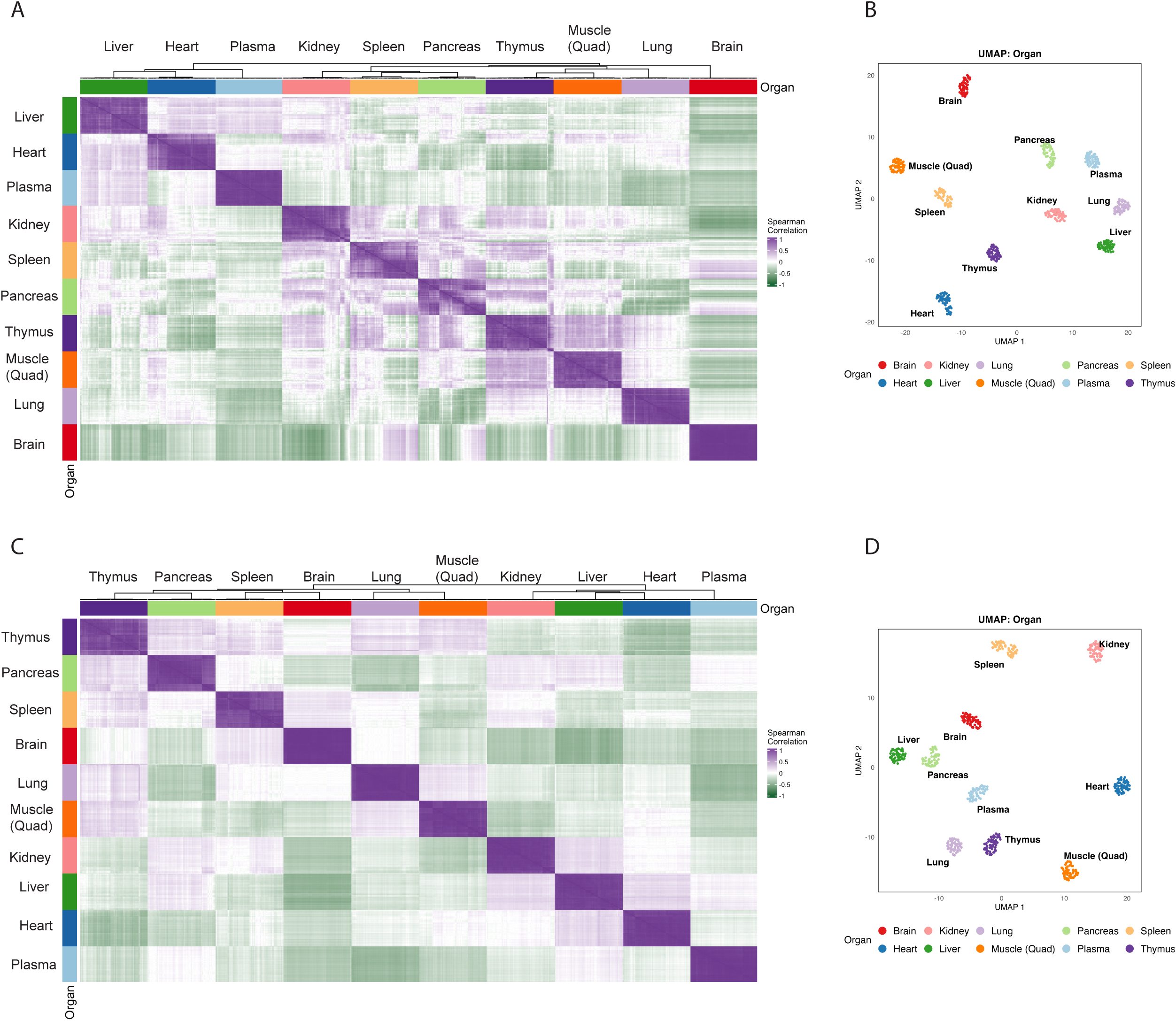
Intra-organ samples have strong correlations in their lipidomes. **a**, Unsupervised clustering of all samples based on the Spearman correlation between each sample. **b**, UMAP of the data labeled by organ. **c**, Unsupervised clustering of all samples based on Spearman correlation excluding TAGs. **d**, UMAP of the data labeled by organ excluding TAGs.

Several organs also showed positive correlations with other tissues, reflecting shared lipid features. The strongest average inter-organ correlation was observed between the thymus and quadriceps muscle (0.32), suggesting similarity in lipid composition between these tissues (Extended Data Fig. 1a). The next highest inter-organ correlation occurred between the pancreas and thymus (0.16). In contrast, the brain exhibited the most negative average inter-organ correlations relative to other tissues. This difference may be partially explained by the lower abundance of TAGs detected in the brain compared with other organs. Plasma samples also showed weak correlations with most tissues, suggesting that contamination of tissue lipid profiles by circulating plasma lipids was minimal. Consistent with these findings, Uniform Manifold Approximation and Projection (UMAP) analysis revealed clear clustering of samples by organ (Fig. 2b), further highlighting strong intra-organ similarity and distinct lipid signatures across tissues.

Because our transition list was enriched for TAGs, we next separated TAGs from non-TAGs in our dataset and asked whether correlations within and between organs were maintained. Organs maintained they separation from each other using both Spearman correlation (Fig. 2c) and UMAP clustering (Fig. 2d). Spearman correlation of non-TAG lipids (Extended Data Fig. 1b) revealed that the brain still had the weakest correlations with other organs, indicating that the lower abundance of TAGs in the brain does not drive differences with other organs, and that these differences must be driven by more abundant lipids in the brain like phospholipids^25^. Removing TAGs reduced the correlation between quadriceps muscle and thymus from 0.32 to 0.16 (Extended Data Fig. 1b), suggesting that TAGs are the main driver of similarity in lipid profiles between the quadriceps muscle and thymus.

Finally, we observed changes in organ clustering when using only TAGs. The brain, lung, liver and plasma maintained intra-organ clustering, but all other others lost their separation from each other (Extended Data Fig. 1c). This suggested substantial variability in triglyceride composition by age and/or sex within those organs, consistent with previous studies showing that triglyceride levels fluctuate depending on physiological and metabolic state^26^. Overall, these results suggest that TAGs drive organ similarities and differences in a subset of organs.

### Aging leads to organ-specific changes in lipid composition

We next determined how individual lipids change with age in each organ and saw that each organ had a distinct pattern of changes in lipids during aging (Fig. 3a). The thymus and quadriceps muscle had the highest percentage of lipids that changed significantly with age, whereas the kidney and lung had the fewest age-associated lipid changes. In the thymus, most significant changes were increases in lipid levels, whereas most changes in quadriceps muscle lipid levels were decreases. (Fig. 3b). Both the thymus and muscle are organs that undergo major remodeling with age, which could help explain the lipid changes observed. The thymus undergoes thymic involution, a gradual shrinking of the thymus characterized by a loss of epithelial tissue that is replaced by adipocytes^27^. Muscle undergoes sarcopenia, which is a gradual loss of muscle mass characterized by reduced muscle strength and function^28^. These processes may be reflected in the lipid remodeling observed in these tissues. The thymus, pancreas, and quadriceps muscle also had large magnitudes of changes in lipid levels, indicating pronounced lipidomic remodeling with age. Whereas the plasma and liver had smaller magnitude changes in lipid levels, despite having a large number of age-associated changes in lipid levels (Fig. 3c). We next reclustered our UMAP analysis based on age (Fig. 3d), which showed age-dependent separations in the quadriceps muscle and lung. Thymus aging was characterized by an increase in levels of TAGs at all ages compared with 1-month-old mice (Fig. 3e).

**Figure 3:**
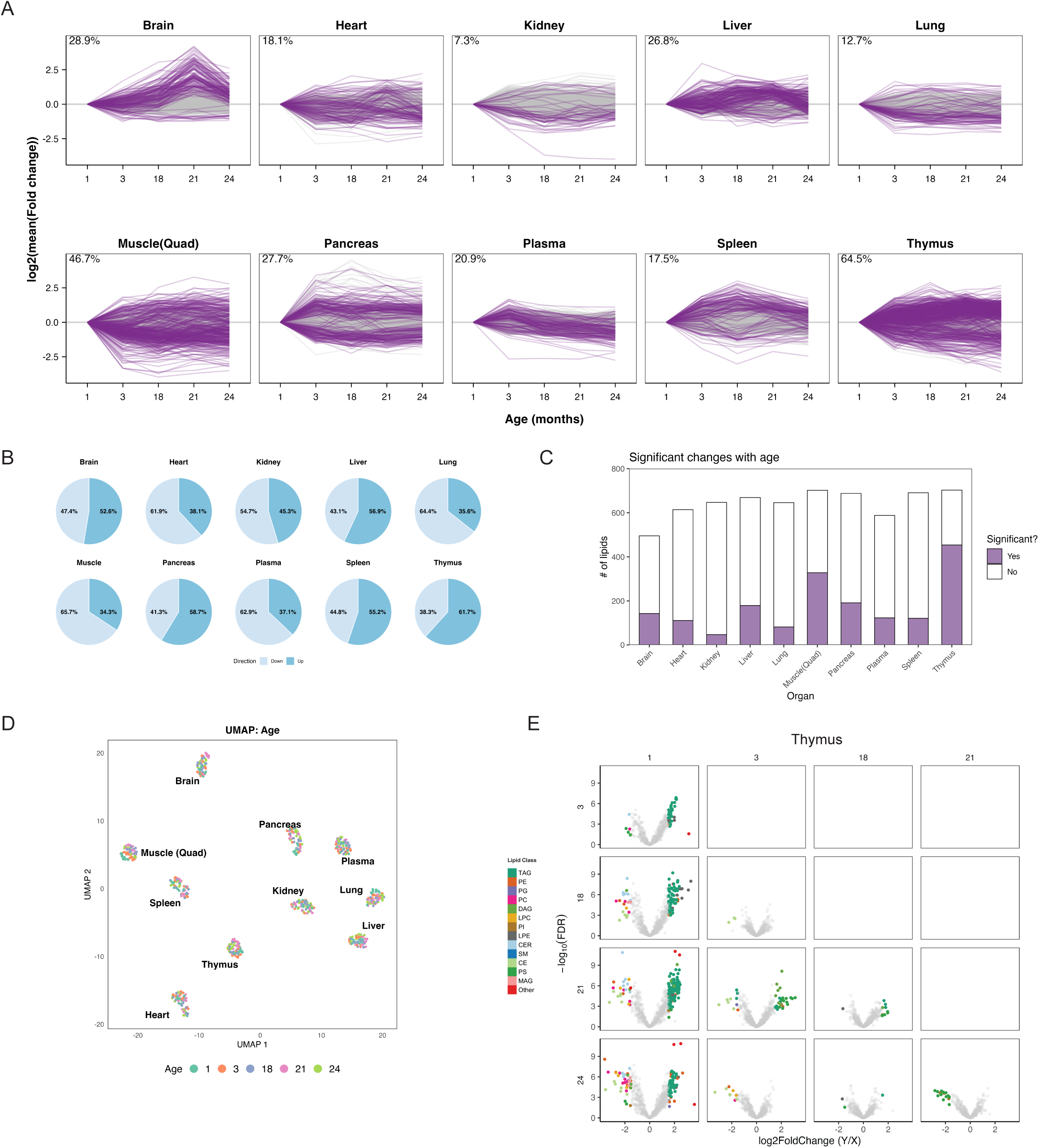
Aging affects each organ differently. **a**, Trajectory of lipid levels in each organ. Each line represents a lipid and the log_2_ fold change relative to the 1-month-old mice. Purple lines represent lipids that show at least one significant change with age (FDR <.05 and mean fold change < -1 or >1), gray lines represent lipids with no significant changes at any age. The percentage of lipids that show at least one significant change is shown for each organ. **b**, Proportion of total lipid comparisons that show either an increase (up) or a decrease (down) compared to 1-month-old mice. **c**, Number of significant lipids per organ. Purple represents significantly changed lipids. **d**, UMAP of the data labelled by age. **e**, Distribution of lipid changes in the thymus. FDR < 0.05, log_2_ mean fold change <− 1 or >1.

We next re-examined the age-dependent lipidome changes by: 1. Excluding TAGs; and 2. Using only TAGs. Organs had similar clustering between non-TAG clustering (Extended Data Fig. 2a) and all lipid clustering (Fig. 3d), further supporting our correlation analyses (Fig. 2a and Fig. 2c). When excluding TAGs, the one-month-old brain samples clustered separately from the older age groups, indicating a strong effect of adolescent developmental on non-TAG lipid composition (Extended Data Fig. 2a).

Age-associated lipid trajectories differed between TAG and non-TAG lipids. The heart, plasma, and brain had fewer age-associated changes in non-TAG lipids (Extended Data Fig. 2b) than in TAGs (Extended Data Fig. 2c), This indicates that TAGs drive age-related lipid variation in these tissues. The quadriceps muscle, pancreas, and thymus maintained age-associated changes in lipid levels both with and without TAGs, consistent with broader lipid remodeling during aging.

### Quadriceps muscle lipidome remodeling during aging is different from cardiac muscle

Quadriceps muscle lipids had the strongest difference between age groups out of all the organs. The one-month-old and three-month-old mice clustered separately from the older age groups (Fig. 4a), with this separation even greater when excluding TAGs (Fig. 4b). We next determined which lipids might drive these age-associated differences and identified distinct clusters of lipids that decreased with age (Fig. 4c), which were enriched for PE and PC lipids (Fig. 4d). The trajectories of the PE and PE lipids in the muscle showed both significant decreases and increases with age (Fig. 4e). The largest age-associated changes occurred among PC and PE lipids containing a 22:4 fatty acyl chain (Fig. 4f), with many of them showing significant decreases with age, including PC(16:0_22:4), PC(18:2_22:4), PE(O-16:0_22:4), PC(18:0_22:4), PE(18:0_22:4), PC(18:1_22:4), PE(18:2_22:4), PE(16:0_22:4) and PE(18:1_22:4) (Fig. 4f). The 22:4 fatty acyl chain is adrenic acid, a polyunsaturated fatty acid synthesized from arachidonic acid. Arachidonic acid is implicated in regulation of metabolism, for example alterations in arachidonic acid is associated with insulin resistance^29^. Arachidonic acid supplementation in mice can also to protect from hyperglycemia-induced muscle atrophy^30^, highlighting a potential link between these lipid pathways and muscle aging. Notably, reductions in PC and PE lipid species have been reported in skeletal muscle from human patients with metabolic syndrome^31^, suggesting that age-related lipidomic changes in muscle may overlap with lipid alterations associated with metabolic dysfunction. Together, these findings demonstrate an age-associated reduction in specific PC and PE species in quadriceps muscle, particularly those containing adrenic acid, and that altered arachidonic acid lipid metabolism may play a role in the muscle aging process.

**Figure 4:**
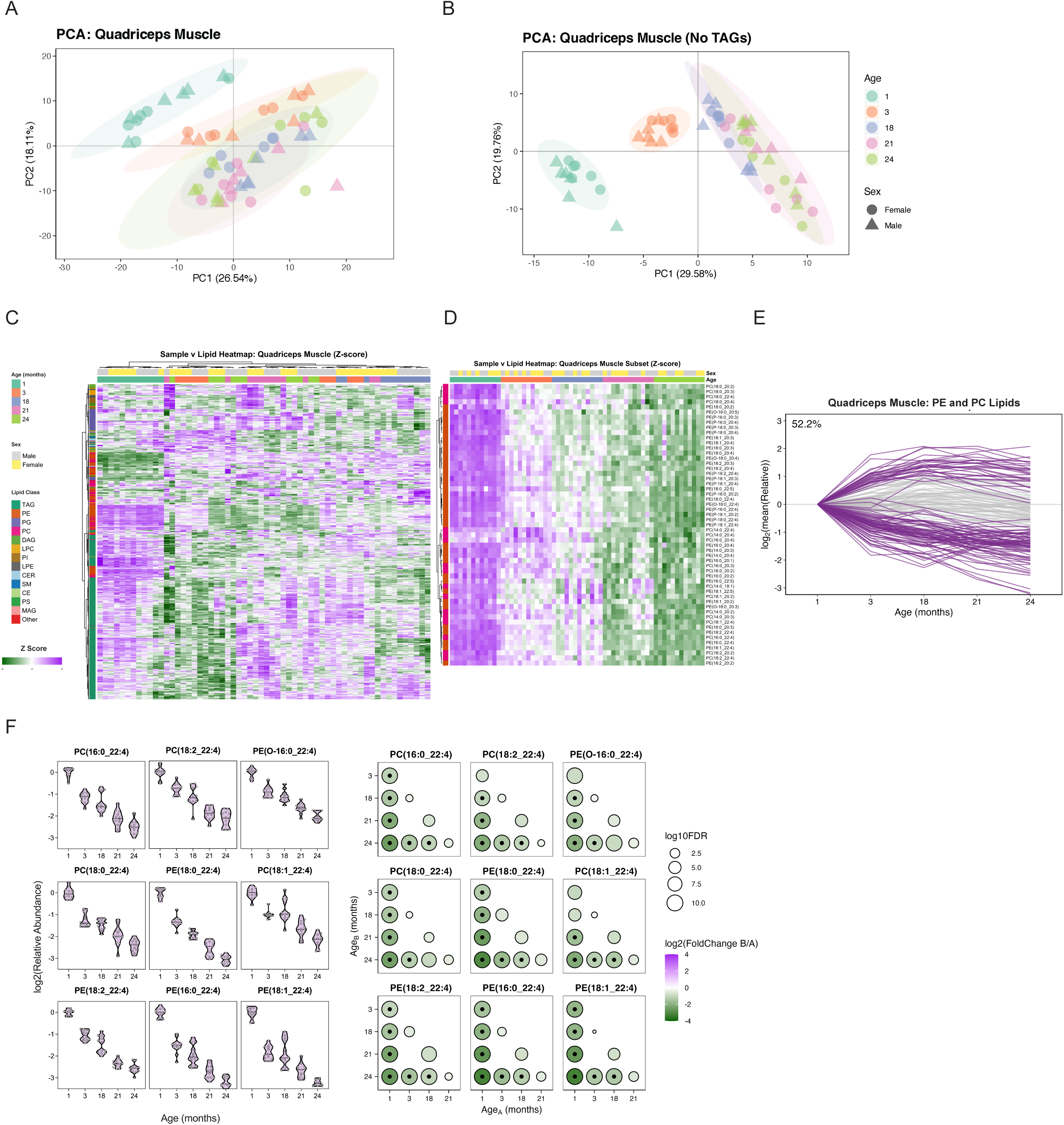
Quadriceps muscle undergoes significant age-associated changes. **a**, PCA of quadriceps muscle samples analyzing all lipids. Sex distinguished by shape, age by color. **b**, PCA of quadriceps muscle samples analyzing non TAGs. Sex distinguished by shape, age by color. **c**, Unsupervised clustering of quadriceps muscle lipid abundance. **d**, Abundance of PCs and PEs = that change with age in the quadriceps muscle. **e**, Trajectory of PEs and PEs with age in quadriceps muscle. Each line represents a lipid and the log_2_ fold change relative to the 1-month-old mice. Purple lines represent lipids that show at least one significant change with age (FDR <.05 and mean fold change < -1 or >1), gray lines represent lipids with no significant changes at any age. The percentage of lipids that show at least one significant change is shown. **f**, (Left) Abundance of selected PCs and PEs in the quadriceps muscle normalized to 1-month-old mice. Horizontal lines, mean abundances (n = 12 per group). (Right) Significance. Circle diameter = log_10_ FDR for age B relative to age A. Circle color = log_2_ transformed fold change. Black dot = FDR < 0.05.

Given the age-associated trends observed in the quadriceps muscle, we next examined whether similar patterns were present in cardiac muscle. The heart did not show age-dependent clustering by PCA when analyzing all lipids (Extended Data Fig. 3a), non-TAG lipids (Extended Data Fig 3b) and TAGs alone (Extended Data Fig. 3c). However, heatmap did show a clustering of the 1-month-old group as seen in the quad muscle (Extended Data Fig. 3d). We also observed the specific PCs and PEs, which had an age-associated decrease in quadriceps muscle (Fig. 4c), showed a similar age-based trend in the heart, although the magnitude of this effect was reduced (Extended Data Fig. 3E). Collectively, these results indicate that while cardiac and quadriceps muscle share certain lipid aging signatures, the age-associated lipidome changes are more pronounced in the quadriceps muscle than in cardiac muscle. This matches the observation that cardiac muscle does not generally undergo age-associated sarcopenia, unlike the quadriceps muscle.

### Sex impacts lipid metabolism

We next identified age-independent sex differences in lipid composition. We ranked the ratios of lipid abundances between males and females for each organ at each age, which revealed sex-specific changes in lipids. Plasma exhibited consistent sex-based differences at all ages, whereas the heart, pancreas, and thymus displayed sex-dependent lipid differences at specific time points (24 months, 21 months, and 1 month, respectively) (Fig. 5a).

**Figure 5:**
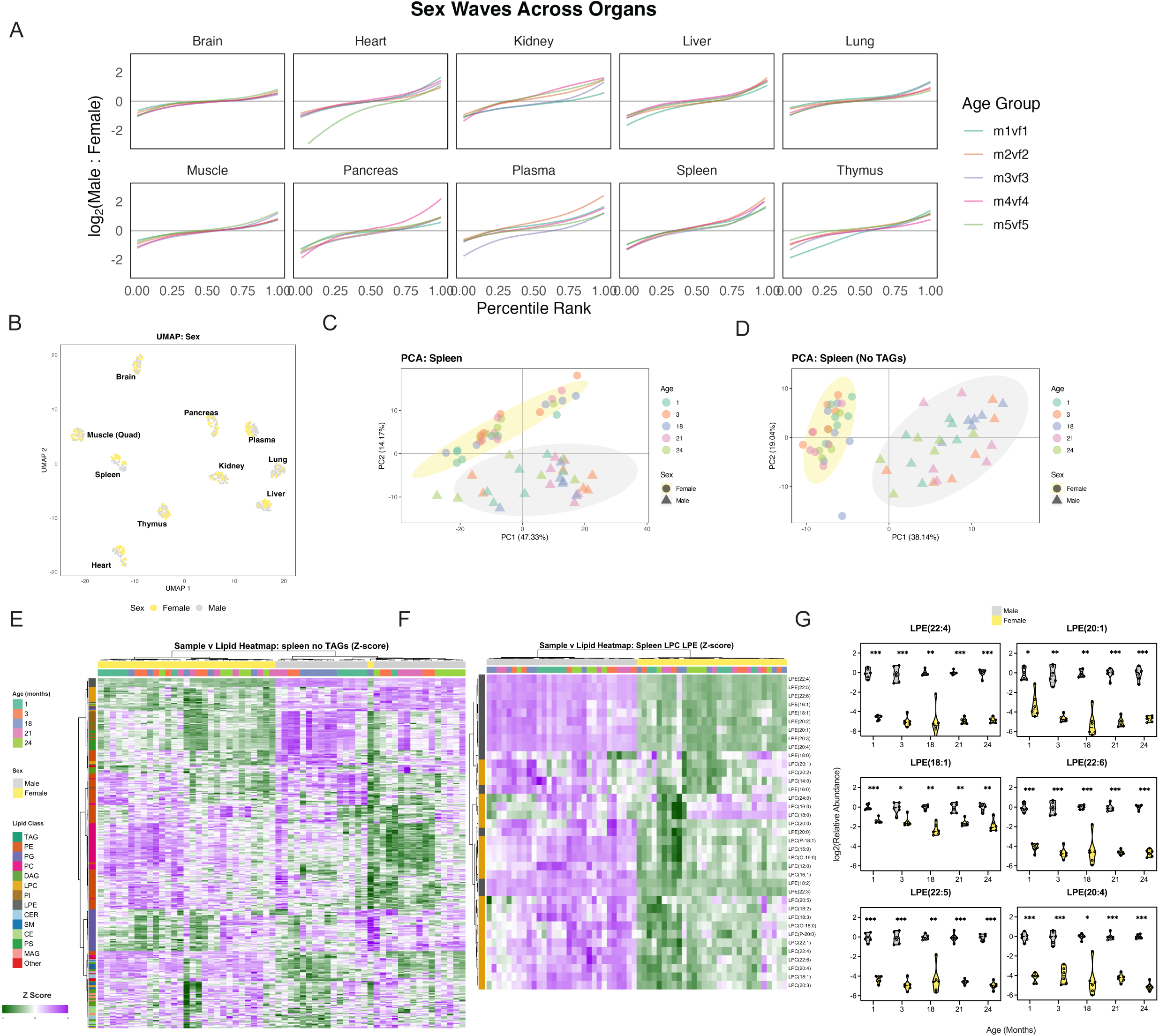
Male and female spleens have distinct lipidomes. **a**, Sex-specific differences in lipids at each age. Each line shows the log_2_ ratio of lipid levels in males to females (color = age). Lipids are ordered by size of the ratio in each organ. **b**, UMAP of data labelled by sex. **c**, PCA of spleen samples analyzing all lipids. Sex distinguished by shape, age by color. **d**, PCA of spleen samples excluding TAGs. Sex distinguished by shape, age by color. **e**, Unsupervised clustering of spleen samples. **f**, Heatmap of spleen LPEs and LPCs. **g**, Abundance of selected LPEs in male and female spleens normalized to 1-month-old male mice. Horizontal lines, mean abundances (n = 12 per group). **FDR < 0.01, ***FDR < 0.001 for male to female comparisons.

Male and female spleens clustered separately using both UMAP (Fig 5b) and PCA (Fig. 5c), The sex difference was even more pronounced when excluding TAGs (Fig. 5d), suggesting that non-TAGs drive these differences. We next identified which lipids could be driving the sex-difference in the spleen (Fig. 5e). The largest difference in lipid levels between male and female spleens were in LPC and LPE lipids, with females having lower levels of both (Fig. 5f). Significant decreases in LPE lipids were identified in females compared to males, including LPE(22:4), LPE(20:1), LPE(20:4), LPE(18:1), LPE(22:6) and LPE(22:5). (Fig. 5g). Given the central role of the spleen in immune function and known sex differences in immune responses and immune aging^32,33^, these findings may reveal a potentially important lipid distinction between male and female mice. Although further work is required to determine whether these lipid-based changes contribute to sex-specific immune phenotypes.

## Discussion

### Thymus and muscle lipid metabolism are strongly affected by age

Our data show that aging affects the lipidome of each organ we investigated, with the kidney having the fewest changes and the thymus and muscle having the largest and most changes. The direction of these changes in the thymus and quadriceps muscle differed, with the thymus having a majority of lipids increase during aging, while the majority decreased in the quadriceps muscle. These shifts in lipid composition may reflect age-related declines in organ function, particularly in the thymus, where our previous work demonstrated age-associated reductions in metabolite levels, including nucleotides.

Thymic involution is characterized by the gradual loss of thymic size and structure and is accompanied by changes in cellular composition, including reduced T cell development. This is consistent with RNA-seq data showing that proliferation-associated gene expression signatures decline with age across multiple immune compartments of the thymus^15^. Thymic involution is also associated with a loss of epithelial cells and an accumulation of adipocytes, which may help explain the global increase in lipid levels observed in the thymus with aging.

### Muscle aging is characterized by a gradual decrease in PC/PE lipids

Our dataset included two different types of muscle: the quadriceps muscle and cardiac muscle. Notably, these two tissues did not show a strong correlation in their lipidome, suggesting differences in overall lipid composition. The quadriceps muscle had the second-highest number of significantly changing lipids with age, whereas the heart had the third-lowest, indicating distinct aging trajectories between skeletal and cardiac muscle.

In the quadriceps, a clear shift was observed after the 1-month age group, with the youngest groups being most similar to each other and markedly distinct from older age groups. Several lipids that changed significantly with age were phosphatidylethanolamine (PE) and phosphatidylcholine (PC) species, particularly those containing adrenic acid (22:4), which gradually decreased with age. In cardiac muscle, decreases in these lipids were also observed, although to a lesser extent. Together, these findings suggest that PC and PE lipids, especially those containing adrenic acid, may play a role in, or reflect, age-associated muscle decline in skeletal muscle and, to a lesser extent, cardiac muscle.

### Male and female spleens have distinct lipidomes

Our study identified sex-associated differences in the lipidome across multiple organs. The organ most strongly influenced by sex was the spleen, which showed clear separation between male and female lipidomes at each age. Differential analysis of these clusters identified enrichment of LPC and LPE lipids in male samples compared to females. This enrichment was particularly pronounced for the LPE class, in which the majority of LPEs were lower in females. Given known sex differences in immune regulation and the central role of the spleen in immune function, these findings suggest that sex-specific differences in LPE abundance may contribute to sex-based differences in immune-related processes.

In summary, our study uncovers age- and sex-dependent changes in lipids in 10 organs. These changes reveal fundamental insights into the aging process and identify new therapeutic targets to maintain organ health.

### Study limitations

Our study uses LC-MS/MS based lipidomics to uncover how lipid pools change during aging in 10 organs in male and female C57BL/6 mice, however using an untargeted technique would reveal how more lipids change with age and sex.

We used only one mouse strain in the study, C57BL/6NCrl, as it is a commonly used strain, which makes our data relevant to many other fields. The strain also matches the strain used in our metabolic atlas of mouse aging and the Tabula Muris Senis, allowing integration of multiple datasets. Although we expect many of our observations to be extendable to multiple species, it will still be necessary to characterize how the lipidome changes during aging in other systems such as non-human primates and humans.

## Methods

### Animals

We purchased male and virgin female C57BL/6NCrl mice from Charles River (RRID:IMSR_CRL:027) and housed them at USC. Mice were housed under identical conditions at both Charles River and USC (12 hr/12 hr light dark cycle) and fed the same diet ad libitum (NIH-31 diet). The 1-month-old, 3-month-old, and 18-month-old mice were kept at USC for one week before harvesting. The 21-month-old and 24-month-old mice arrived aged 18 months and were aged at USC for the remaining 3 months and 6 months respectively.

### Organ harvest

All tissues were collected beginning at 3 pm on each day of harvest. We used 4% isoflurane to anaesthetize the mice. We then weighed the mice and collected blood via cardiac puncture into EDTA collection tubes and placed them on ice. We then collected in the following order: heart, thymus, lungs, liver, pancreas, spleen, kidney, quadriceps muscle, and brain. The organs were immediately snap frozen in liquid nitrogen and stored at -80°C until metabolite extraction. Organ collection finished by 5 pm. All animal care and procedures were approved by the USC Department of Animal Research. To obtain plasma, we centrifuged EDTAtubes containing blood at 4°C at 2,000 g for 15 minutes. We collected supernatants and stored at -80°C until further processing.

### Lipid extraction

For plasma samples, we added 20 μL of plasma to 80 μL 100% LC-grade methanol. For tissue homogenization, we put tissue samples in 10 μL ice-cold 5:1 LC-grade Methanol:Ultrapure-water per mg tissue and homogenized them using a bead homogenizer (Bead Mill 4, Fisherbrand). For all samples, we next added ice cold methyl tertiary-butyl ether (MTBE) to each sample and placed on an orbital shaker for 1hr at room temperature. Phase separation was induced by adding Ultrapure water, so the final concentrations of MTBE/MeOH/H2O were 10/3/2.5. We centrifuged plasma supernatants and tissue homogenates at 4°C at 21,300 g for 10 minutes and collected the lipid containing top phase. We dried down 300 μL of supernatant from each sample (equivalent to 10 mg of tissue) using a vacuum centrifuge and stored them at -80°C until further processing.

### Sample preparation

All organs except brain: 100 µL of 1:2 IPA:ACN was added to the dried extracts, vortexed for 10 seconds, then reconstituted on an orbital shaker for 10 min at 2000 rpm. Samples were then centrifuged for 5 min at 2°C and the supernatant aliquoted for LC-MS injection.

Brain: 100 µL of 60:30:4.5 MTBE:MeOH:H2O was added to the dried extracts, and underwent the same reconstitution and centrifugation procedure above. The resulting supernatant was diluted 4-fold in 1:2 IPA:ACN and aliquoted for LC-MS injection.

### Liquid chromatography-tandem mass spectrometry (LC-MS/MS)

Lipidomic analyses were performed using a targeted LC–MS/MS approach. Samples were separated by hydrophilic interaction liquid chromatography (HILIC) based on a previously described protocol (ref). A 10 µL aliquot of each sample was injected onto an XBridge Amide column (4.6 × 150 mm, 3.5 µm; Waters Corporation, Milford, MA) maintained at 35°C using a 1290 Infinity II HPLC system (Agilent Technologies). Mobile phase A consisted of 10 mM ammonium acetate in 95:5 acetonitrile:water, and mobile phase B consisted of 10 mM ammonium acetate in 50:50 acetonitrile:water, with the pH adjusted to 8.4 using ammonium hydroxide. The gradient (min/%B) was as follows: 0/0.1, 6/6, 10/25, 11/98, 13/100, 18.6/100, 18.7/0.1, and 28/0.1. The flow rate was set to 0.7 mL/min from 0–13.4 min, increased to 1.5 mL/min from 13.5–23 min, and returned to 0.7 mL/min from 23.5–28 min.

The column effluent was directed to an OptiFlow Turbo V electrospray ionization source coupled to a QTRAP 6500+ mass spectrometer (AB Sciex LLC). Data were acquired in targeted multiple reaction monitoring (MRM) mode in both positive and negative ionization polarities. Lipid species were identified and quantified based on detection of characteristic fragment ions at defined retention times, ensuring specificity in complex biological matrices.

For positive ion mode, source parameters were as follows: Curtain Gas (CUR) 35 psi, Collision Gas (CAD) medium, IonSpray Voltage (IS) 4500 V, Temperature (TEM) 550°C, and Ion Source Gases 1 and 2 (GS1, GS2) set to 30 and 40 psi, respectively. Negative ion mode parameters were identical, except the IonSpray Voltage was set to −4500 V.

### Data Analysis

Raw lipidomics data was imported into Skyline (MacCoss Lab Software) and checked for quality. 4 external standard injections were made before the 1^st^, 20^th^, 40^th^, and after the 60^th^ sample of the sequence respectively using 10 µL of 5 ng/µL concentration in 1:2 IPA:ACN of EquiSPLASH™ LIPIDOMIX® Quantitative Mass Spec Internal Standard (Avanti Polar Lipids Inc.), showing retention times and signal were consistent across the sequence. 8 injections were made of a pooled sample mix in sets of 2 to monitor detector response of individual lipid species throughout the run.

Chromatograms were processed in Skyline Daily (Version 23), where all peaks were integrated and manually reviewed to ensure consistent retention times and proper alignment of fragment ions. In R (v4.3.1), peak areas were normalized to the total ion count for each sample and log₂-transformed. Pareto scaling was applied for principal component analysis. Differential abundance analyses was performed using the *limma* package to calculate fold changes and p-values for age and sex comparisons, with p-values adjusted by the false discovery rate method. All visualizations were generated with *ggplot2* and *pheatmap*.

## Author contributions

Conceptualization, P.J.M.; Methodology, E.E., S.E.P., D.L., G.W., W.C. and P.J.M.; Formal Analysis, E.E., S.E.P. and S.S.; Investigation, E.E., S.E.P., X.S., P.I., S.S., R.M., S.B.K., D.M., A.J.C. and P.J.M.; Resources, P.J.M.; Writing – Original Draft, E.E. and P.J.M.; Writing – Review and Editing, E.E. and P.J.M. with input from all authors; Visualization, E.E., S.E.P. and X.S.; Supervision, R.H.S., W.C. and P.J.M.; Project Administration, P.J.M.; Funding Acquisition, P.J.M.

## Acknowledgements

This work is supported by Impetus Grants 014746-00001; A.J.C. is supported by the National Institute on Aging T32AG052374; R.H.S. is supported by R00AG065200 and R01AG079806 from the National Institute on Aging and the Glenn Foundation for Medical Research and AFAR Grant for Junior Faculty Award; P.J.M. is supported by Impetus Grants 014746-00001, the American Lung Foundation COVID-920798, NIH RECOVER ROA-OTA-21-015J and the Donald E. and Julia B. Baxter Foundation.

## Figure Legends

**Extended Data Figure 1: Correlations in TAGs and non-TAG lipid groups**

**a**, The mean Spearman correlation for every organ comparison in Figure 2a.

**b**, The mean Spearman correlation for every organ comparison in Figure 2c.

**c**, Unsupervised clustering of TAGs in all samples based on Spearman correlation.

**Extended Data Figure 2: Age-associated changes in organ TAG and non-TAG lipids**

**a**, UMAP of the data labeled by age excluding TAGs.

**b**, Trajectory of non-TAG levels in each organ. Each line represents a non-TAG lipid and the log_2_ fold change relative to the 1-month-old mice. Purple lines represent non-TAGs that show at least one significant change with age (FDR <.05 and mean fold change < -1 or >1), gray lines represent non-TAGs with no significant changes at any age. The percentage of non-TAGs that show at least one significant change is shown for each organ.

**c**, Trajectory of TAG levels in each organ. Each line represents a TAG and the log_2_ fold change relative to the 1-month-old mice. Purple lines represent TAGs that show at least one significant change with age (FDR <.05 and mean fold change < -1 or >1), gray lines represent TAGs with no significant changes at any age. The percentage of TAGs that show at least one significant change is shown for each organ.

**Extended Data Figure 3: Age-associated changes in heart lipids**

**a**, PCA of heart samples analyzing all lipids. Sex distinguished by shape, age by color.

**b**, PCA of heart samples excluding TAGs. Sex distinguished by shape, age by color.

**c**, PCA of heart samples using only TAGs. Sex distinguished by shape, age by color.

**d**, Unsupervised clustering of heart lipid abundance.

**e**, Abundance of PCs and PEs in the heart (PCs and Pes shown are the same as in Figure 4c).

