## Supplementary figures and images for "The aging mouse lipidome"

### Extended Data Figure 1

Extended Data Figure 1

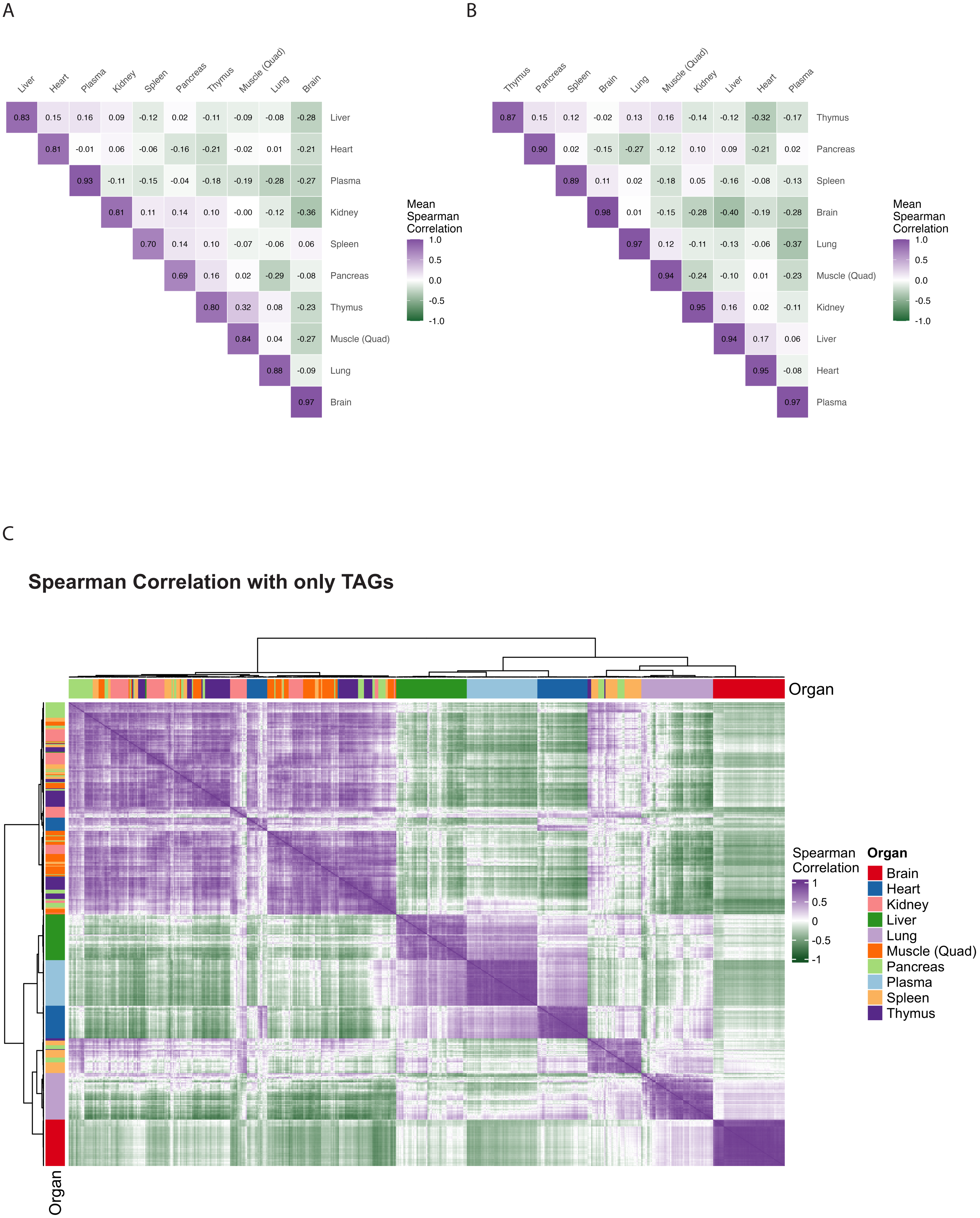

### Extended Data Figure 2

Extended Data Figure 2

A

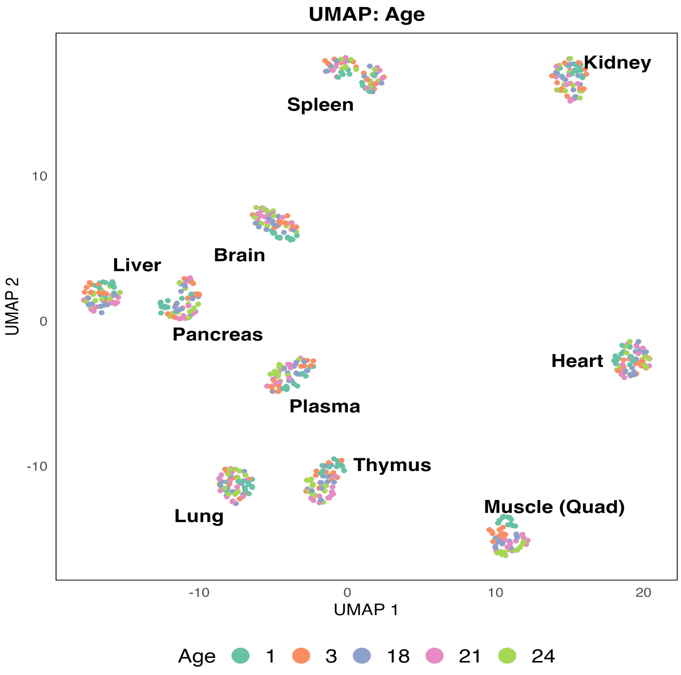


B

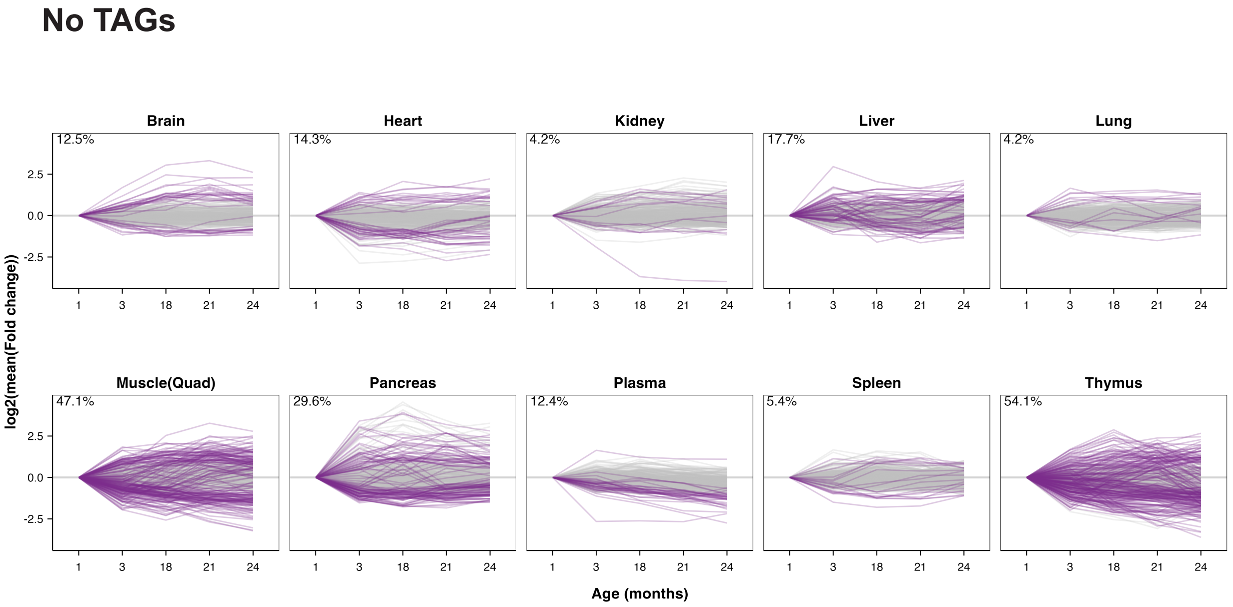


C

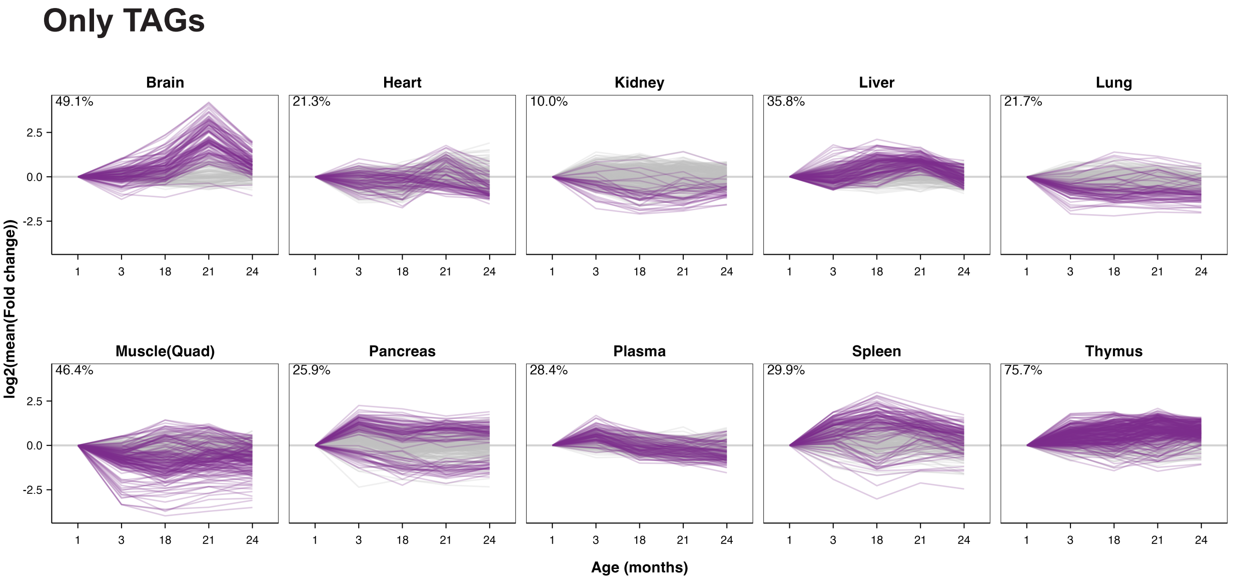

### Extended Data Figure 3

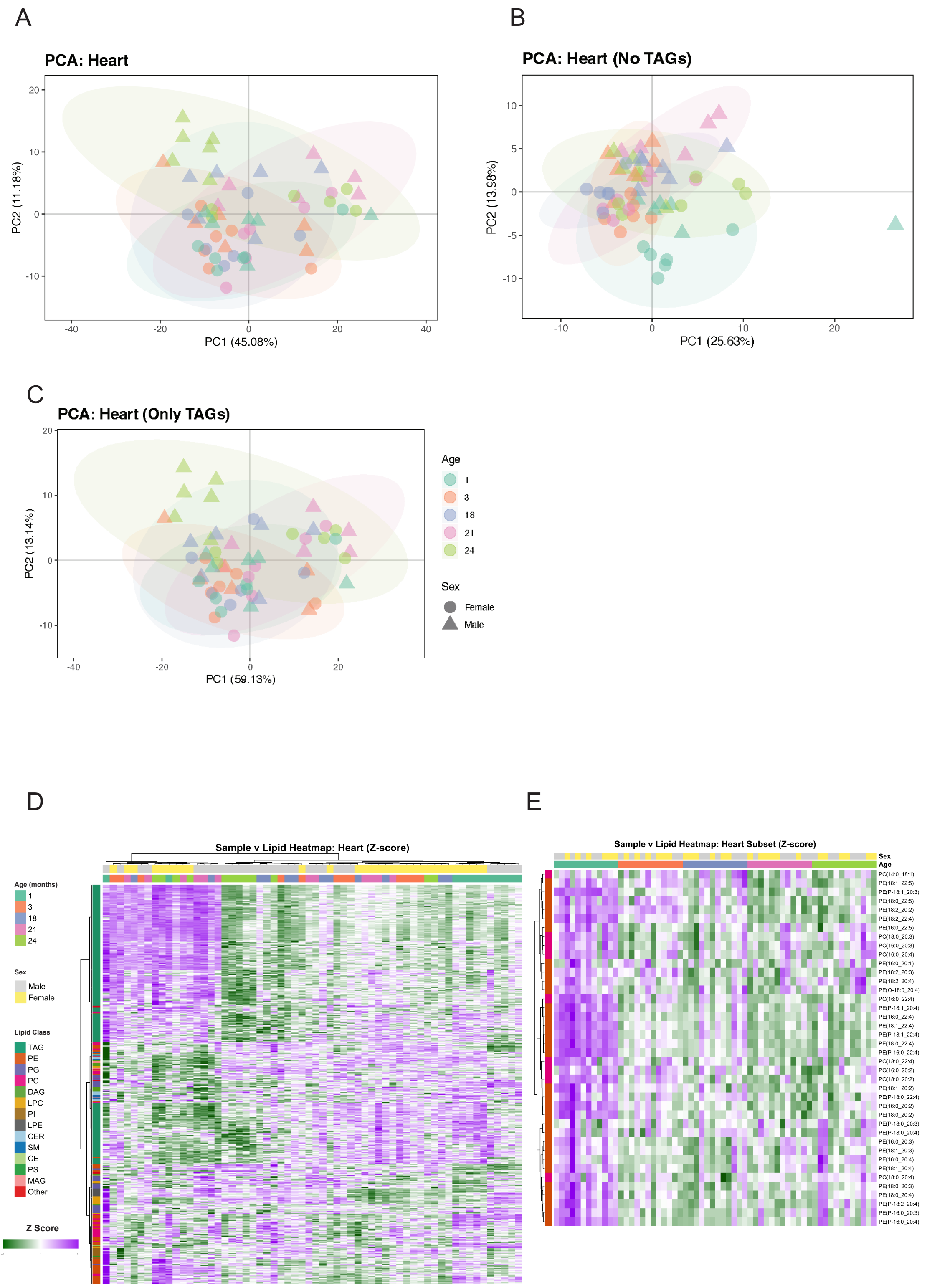
